# Effects of chronic spinal cord injury on relationships among ion channel and receptor mRNAs in mouse lumbar spinal cord

**DOI:** 10.1101/357665

**Authors:** Virginia B. Garcia, Matthew D. Abbinanti, Ronald M. Harris-Warrick, David J. Schulz

**Affiliations:** Division of Biological Sciences, University of Missouri, Columbia, MO USA 65211; Department of Neurobiology and Behavior, Cornell University, Ithaca NY 14853, USA

**Keywords:** Keywords Ion Channels, Receptors, Gene Co-Expression Network, Correlation Analysis

## Abstract

Spinal cord injury (SCI) causes widespread changes in gene expression of the spinal cord, even in the undamaged spinal cord below the level of the lesion. Less is known about changes in the correlated expression of genes after SCI. We investigated gene co-expression networks among voltage-gated ion channel and neurotransmitter receptor mRNA levels using quantitative RT-PCR in longitudinal slices of the mouse lumbar spinal cord in control and chronic SCI animals. These longitudinal slices were made from the ventral surface of the cord, thus forming slices relatively enriched in motor neurons or interneurons. We performed absolute quantitation of mRNA copy number for 50 ion channel or receptor transcripts from each sample, and used multiple correlation analyses to detect patterns in correlated mRNA levels across all pairs of genes. The majority of channels and receptors changed in expression as a result of chronic SCI, but did so differently across slice levels. Furthermore, motor neuron enriched slices experienced an overall loss of correlated channel and receptor expression, while interneuron slices showed a dramatic increase in the number of positively correlated transcripts. These correlation profiles suggest that spinal cord injury induces distinct changes across cell types in the organization of gene co-expression networks for ion channels and transmitter receptors.

## INTRODUCTION

Chronic spinal cord injury (SCI) leads to a variety of neurological complications across sensory, motor, and autonomic systems (Wienecke et al., 2010; Chen et al., 2013; Lee-Liu et al., 2014). An initial state of depressed motor system activity in acute injury often gives rise to chronic changes in motor network excitability that can cause complications such as spasticity in the affected limbs (Little et al., 1999; Dennis et al., 2003; Hultborn, 2003; Frigon and Rossignol, 2006; Nielsen et al., 2007; Rossignol et al., 2007). Additionally, hyperexcitability below the site of the injury can result in chronic pain (Carlton et al., 2009; Hulsebosch et al., 2009; Finnerup et al., 2014; Widerström-Noga, 2017), and influence autonomic functions that can give rise to dangerous complications such as autonomic dysreflexia (Hagen et al., 2011; Bauman et al., 2012; de Groat and Yoshimura, 2012). The varied impacts of injury on the networks below the affected spinal level strongly suggest that the loss of descending inputs can dramatically alter systems and networks that were not themselves damaged by the injury. The chronic nature of these effects reflects long-term changes that have been shown to extend to the level of gene expression and transcriptional regulation. It is reasonable to hypothesize that changes in gene expression associated with chronic SCI will vary based on the region of the cord analyzed. Therefore, a more complete understanding of the impacts of SCI requires an understanding of these system-specific changes. In addition, it is important to study how patterns of gene co-expression change across genes and gene families (Ryge et al., 2010; Garcia et al., 2014), as neuronal output is ultimately the result of the relative levels of expression of ion channels and neurotransmitter receptors which cooperate to shape their electrophysiological activity.

It is clear that spinal cord injury causes widespread changes in gene expression of the spinal cord across numerous gene families; this has been demonstrated by various approaches including qPCR (Esmaeili and Zaker, 2011; Di Narzo et al., 2015), microarray analyses (Carmel et al., 2001; Ryge et al., 2008, 2010; Wienecke et al., 2010; Liu et al., 2014) and RNAseq (Chen et al., 2013; Lee-Liu et al., 2014). There is also growing appreciation that in addition to overall levels of transcript, expression levels of some genes vary in parallel over repeated measurements: this co-regulation of functionally interacting channels and receptors may be essential to maintain appropriate neuronal output (Amendola et al., 2012). For example, motor neurons of crustaceans express a common set of voltage-dependent channel subunits, but the correlated levels of these transcripts differ across identified cell types, and the ratio of co-expression can also differ (Schulz et al., 2007; Tobin et al., 2009). Furthermore, these expression relationships are not fixed: they are both activity-dependent and influenced by neuromodulator release from descending inputs (Temporal et al., 2012, 2014; O’Leary et al., 2013). These correlated channel and receptor expression patterns, as well as their lability, are also detectable in mammalian spinal cord: different spinal cord levels exhibited distinct patterns of correlated channel and receptor expression in adult mice, and change over post-natal development (Garcia et al., 2014). In this study we bring this emerging perspective on correlated expression patterns to the effects of SCI.

It is currently impossible to gain a consensus view of injury-induced changes in gene expression, as studies vary widely in their model systems from rodents to human, injury type and cord level, and sample collection methods which range from single neurons to whole spinal cord. This study does not attempt to reach a definitive conclusion regarding changes in gene expression in the cord after injury. Rather, we seek to shed light on three distinct aspects of gene expression associated with SCI. We do so by measuring the mRNA abundance for channels and receptors in the lumbar spinal cord following a complete thoracic level transection. First, our results demonstrate that SCI does not result in uniform changes in gene expression across different longitudinal sections of the lumbar spinal cord, which are relatively enriched for different cell classes. Second, changes in gene expression after SCI do not simply alter the abundance of a given gene transcript, but also substantially change relative co-expression relationships *across* genes, measured as the correlation in expression between all possible pairs of measured genes. Third, changes in gene co-expression networks after SCI vary profoundly in different regions of the spinal cord. Taken together, our results shed new light on the complex relationships in gene expression in the spinal cord, and the differential effects of injury on these relationships both across neuron types and gene families.

## METHODS

### Animals

All animal protocols were approved by the Cornell University Institutional Animal Care and Use Committee and in accordance with guidelines from the National Institutes of Health. For SCI surgery, P28-35 C57BL/6 ChAT^BAC^-eGFP mice (Jackson laboratories, Stock No 007902, B6.Cg-Tg(RP23-268L19-EGFP)2Mik/J) were anesthetized with intraperitoneal (I.P.) ketamine (100 mg/kg) and xylazine (5 mg/kg). Subcutaneous 0.06 mg/kg Buprenorphine was administered at the start of surgery. Dorsal skin was shaved, depilated, and sterilized with alternating swabs of 70% ethanol and betadine. Using aseptic technique, a skin incision was made above the thoracic spine and the musculature was teased apart to expose the spinal column. The spinal cord was exposed by gently stretching the T8 and T9 vertebrae apart. Using iris scissors and fine forceps, the spinal cord was completely transected at T8-T9. A sterile tuberculin needle with a slightly hooked tip was passed across the transection multiple times to confirm complete transection. The incision was closed with tissue glue. Subcutaneous ketoprofen (2 mg/kg) and lactated ringers (0.5 cc/20 g body weight) were administered immediately. Injured mice received buprenorphine twice daily for two days and ketoprofen daily for four days post-SCI. Bladders were manually expressed until normal bladder function returned. A total of 60 animals were used in these studies.

Recovery of function after this complete lesion was limited. Mice were scored twice weekly on a 5-point scale to monitor hindlimb spontaneous movement, with 1 representing no movement and 5 representing normal left-right alternation and full body support. A second measurement monitored the motor response to a rapid tail pinch, with 1 representing uncoordinated hindlimb movement ant 5 representing activation of a full locomotor response with proper left-right alternation. For both measures, hindlimb movement was almost completely absent in the first week, and gradually improved to a plateau at 3 weeks after injury. At this plateau, mice showed occasional spontaneous movement of the hindlimbs with partial lifting of the body, without left-right alternation. The response to tail pinch improved more, with marked hyperextension and left-right alternation of the limbs and partial lifting of the body weight. There was no further improvement with time after 3 weeks.

Reductions in spinal cord diameter have been reported after SCI, especially near the site of the lesion (Freund et al., 2013; Hooshmand et al., 2014). Because the slicing procedure used to collect samples for this study (described below) requires relatively consistent anatomy among control and injured animals, individuals that exhibited a detectable reduction in cord size in the lumbar region (L1-L5) after the T8-T9 lesion were eliminated from the analysis. Animals that exhibited significant shrinking cord size after injury often presented with unnatural hindlimb positioning at rest, including foot pads that pointed largely upwards, with no improvement in motor activity as described above. These changes were grounds for exclusion from the experiment.

### Slicing procedure

The slicing procedure is summarized in Figure 1. At two-three months of age (and 1-2 months post-SCI for injured animals), mice were deeply anesthetized by I.P. ketamine (150 mg/kg) and xylazine (15 mg/kg). The lumbar spinal cord was rapidly dissected and meninges removed in ice-cold oxygenated (95/5% O_2_/CO_2_) glycerol-based aCSF ([in mM] 222 glycerol, 3 KCl, 1.18 KH_2_PO_4_, 1.25 MgSO_4_, 2.5 CaCl_2_, 25 NaHCO_3_, and 11 D-glucose). The cord was embedded in 1.5% agarose in glycerol-based aCSF. After rapid cooling, the agarose was trimmed and the embedded cord was mounted on the vibratome (Microm HM650V, Thermo Scientific), and 300 μm longitudinal slices were prepared. The first slice contained ventral white matter and was not used. The next 300 μm slice was then collected; we confirmed these slices contained motor neurons in multiple instances by visualization of ChAT+ motor columns. These slices (referred to as MNslices) contained only lamina 8 and 9. The next 300 μm slice was considered an interneuron-enriched slice (INslices) and collected as such. As a way of validating our method, in multiple experiments, we confirmed this slice contained few or no motor neurons by noting the lack of ChAT+ fluorescence. INslices contained largely neurons from lamina 7, but a small number of neurons from lamina 10 might have also been included. There were no neurons from lamina 6 or any more dorsal portion of the cord in these samples. For whole slice analysis, slices were immediately placed in Trizol (Invitrogen, Carlsbad, CA) for 30 minutes at room temperature, then placed at −80°C until analysis. We used N = 10 of each slice type for subsequent molecular analyses.

**Figure 1.**
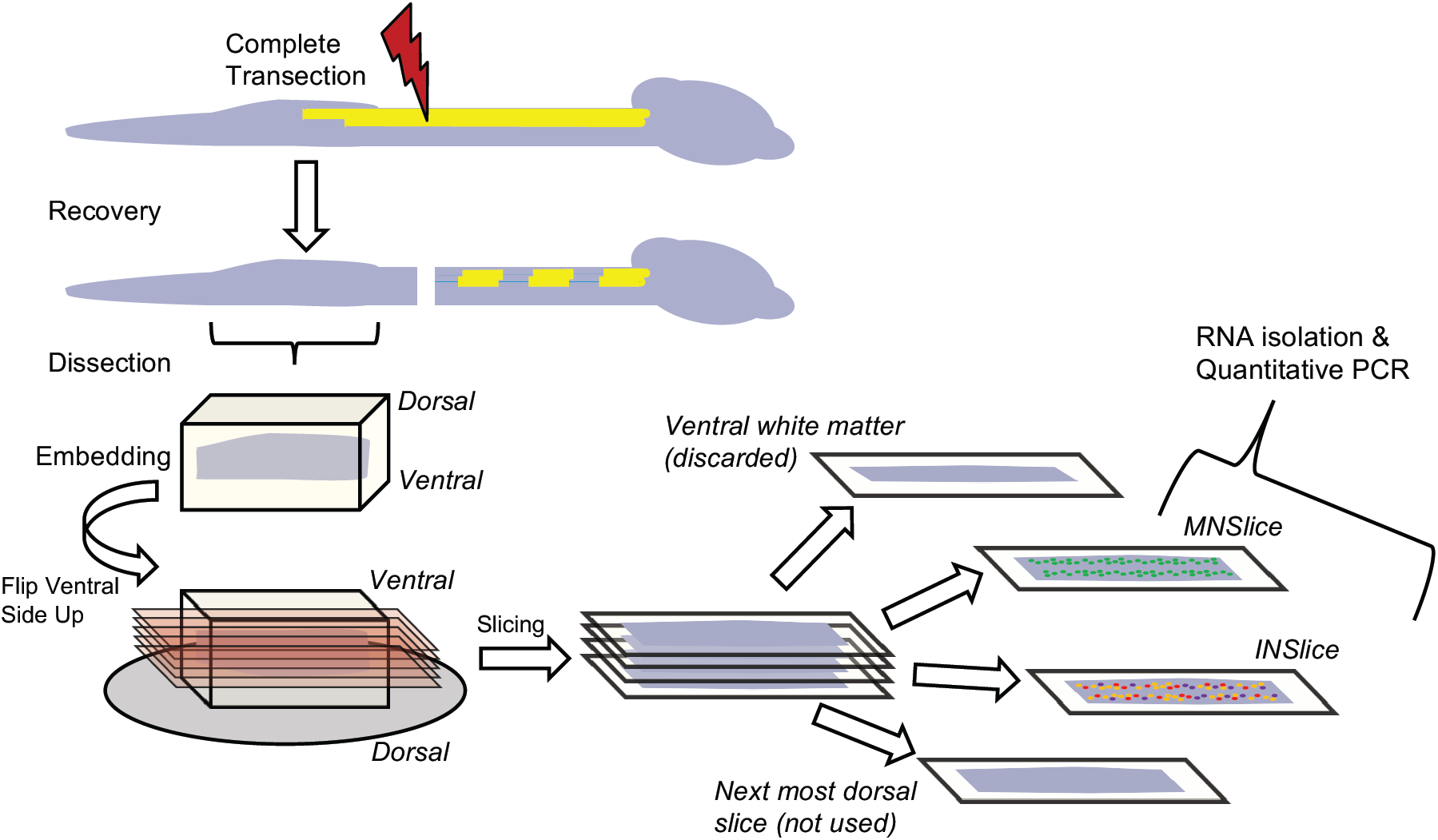
Schematic illustration representing the spinal cord slice procedure used to collect tissues for analysis. Following complete transection at cord level T8-T9, animals were allowed to recover from injury for a period of at least 28 days. Animals were sacrificed, the cord below the injury removed, and embedded for slicing. Longitudinal slices were made from the ventral surface of the cord and progressed in 300 μm increments. Slices of gray matter that contained concentrations of ChAT:GFP labeling were collected as motor neuron enriched slices (MNSlice). After removal of GFP containing neurons, the next most 300 μm dorsal slice was collected as interneuron enriched (INSlice). These slices were immediately placed in TriZol reagent and homogenized for total RNA extraction.

### Individual neuron isolation

For harvesting of individual neurons, slices were transferred to 37°C oxygenated aCSF ([in mM] 111 NaCl, 3 KCl, 1.18 KH_2_PO_4,_ 1.25 MgSO_4_, 2.5 CaCl_2_, 25 NaHCO_3_, and 11 D-glucose) and allowed to cool to room temperature over 45 minutes. GFP+ neurons were visualized under epifluorescent illumination on a Zeiss Axioskop 2 microsope. Fluorescent ChAT:GFP+ neurons located in the motor neuron columns were individually aspirated via a patch-pipette containing standard patch solution made with RNAase-free water. One to eleven neurons (of known quantity for each sample) were aspirated into a single patch pipette. The pipette tip was then broken and the solution aspirated directly into RNA lysis buffer (Zymo Research, Irvine, CA) and placed at −80°C until analysis.

### Quantitative PCR

Total RNA was isolated from spinal cord slices using Trizol according to the protocol provided by the manufacturer (Invitrogen). cDNA was generated from 500 ng total RNA primed with a mixture of oligo-dT and random hexamers that was reverse transcribed in a 20 μl reaction containing a final concentration of 2.5 ng/μl random hexamers, 2.5 μM oligo-dT, 40 U of RNAseOUT RNase inhibitor, and 200 U of SuperScript III reverse transcriptase. Following heat inactivation of the enzyme, samples were diluted 5X in ultrapure water (final volume 100 μl) and used as template in qPCR analyses. From each cDNA pool generated from 500 ng of total RNA, we quantified at least 15 different gene products.

We designed or modified, and independently validated primer sets for use in absolute quantitation of copy number for 50 distinct genes of interest from the mouse (see Appendix A). Eight of these transcripts were not detected in appreciable levels and subsequently removed from the analysis (*GLRA4, SERT, KCNA4, KCNA5, KCNC1, KCNC2, KCND3, KCNQ1*). Primer sets were either modified from previously validated primer sets as listed in PrimerBank (Spandidos et al., 2010), or designed *de novo* using Primer3 software (Untergasser et al., 2012). Primer sets are listed in Appendix A. Detailed methods regarding the primer validation procedures can be found in our previous work (Garcia et al., 2014). qPCR reactions consisted of primer pairs at a final concentration of 2.5 μM, cDNA template, and SYBR master mix (BioRad) according to the manufacturer’s instructions. Reactions were carried out on a CFXConnect (BioRad) machine with a three-step cycle of 95°C-15s, 58°C-20s, 72°C-20s, followed by a melt curve ramp from 65°C to 95°C. Data were acquired during the 72°C step, and every 0.5°C of the melt curve. All reactions were run in triplicate, and the average Ct (cycle threshold) was used for interpolation with the standard curve to generate copy number for a given reaction. Standard curves for each gene were generated from a known copy number of a plasmid containing a partial ORF for a given gene of interest. Plasmids of known copy number were diluted from 10^6^ copies/μl to 10 copies/μl by factors of 10×, and run in identical qPCR reactions to the samples. A line was fit to the standard curve and the subsequent equation for the fit line used to interpolate copy number from Ct values obtained from biological samples.

The unit we use to express all of the qPCR data in this study is “normalized copy number per 500 ng total RNA.” All of the data were normalized relative to Glyceraldehyde 3-phosphate dehydrogenase (GAPDH) expression (Suzuki et al., 2000). Instead of utilizing a relative quantitation method (Pfaffl, 2001), we created a “normalization factor” using GAPDH Ct values (Garcia et al., 2014). Briefly, each sample is normalized to the population mean for each gene of interest by creating an adjustment factor based on GAPDH expression above or below the population average. A normalization factor using GAPDH for a sample *x* was calculated by the following formula:

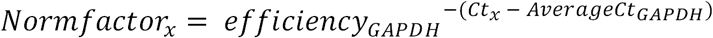

For our assay, GAPDH amplification efficiency was virtually 100%, so this can be simplified as:

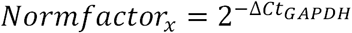

where Δ*CT*_*GAPDH*_ is the difference between the sample GAPDH and the population average, and 2 is the base of the exponential since 100% efficiency results in a doubling of product in each cycle of PCR. This resulted in the normalized copy number for a sample *x* calculated as:

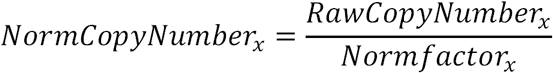

This normalization factor allows us to preserve the overall magnitude of the raw copy number, allowing for comparisons of mRNA abundance *across* genes. In addition, the normalization adjusts the copy number across different slices to account for differences in mRNA abundance that may occur for technical reasons, including variability in RNA extraction and reverse transcription efficiencies. During evaluation of GAPDH expression, two SCI Interneuron slices were found to have very low expression, likely reflecting poor initial RNA quality or extraction, and were eliminated from the analysis. We also determined that for one Control Interneuron sample, 9 of the measured transcripts had poor quality amplification and were not suitable for quantitation; thus this sample was removed from the analysis rather than using only selected data from it. Thus our final sample sizes for analyses are: Control Motor (N = 10), Control Interneuron (N = 9), SCI Motor (N = 10), and SCI Interneuron (N = 8).

### Experimental Design and Statistical Analysis

Overall differences in mean mRNA copy number across groups were tested for normality via the Shapiro-Wilk test, and tested for equal variances via F-test of equality of variances (SigmaPlot v11). Those that were found to violate normality/equal variance were log-transformed and re-tested. Data were then analyzed for each gene separately via two-way ANOVA (SigmaPlot v11) with slice level and injury state as the two factors. Significant ANOVA results were further tested with post-hoc *t*-tests for significant differences between control and injured groups for a given slice level, with multiple comparisons procedures adjusted based on the Holm-Šídák method.

The combined dataset for different slice levels and injured vs. control animals were subjected to hierarchical clustering analysis (SPSS v20) to test for homogeneous clusters among the sample groups. Clustering and construction of dendrograms were generated using Ward’s hierarchical clustering/linkage methods.

Correlation matrices, R-values, and correlograms were generated using Pearson’s tests for every pairwise combination of mRNA levels for all genes that generated detectable expression (R statistics package). For 42 genes this created 861 pairwise comparisons for each experimental group. From the R-values for each of these comparisons, a heat-mapped correlogram was produced where the X- and Y-axes correspond to the pairwise gene comparisons and the correlation coefficient (R-value) from the pairwise Pearson’s tests is represented as a color mapped to a scale from −1 to 1. To account for the statistical error associated with making so many individual comparisons, we performed multivariate permutation tests of correlations for all data in the matrix to estimate the true P-value for each comparison. Each pairwise comparison was subjected to 10,000 permutations to generate an exact P-value [MPTCorr.r, R statistics package, (Yoder et al., 2004; R Development Core Team, 2008)]. In addition, cumulative distribution functions for R-values between two groups were compared by two-sample Kolmogorov-Smirnov tests. Gene co-expression network and differential correlation analyses were performed with the DiffCorr R package (Fukushima, 2013).

## RESULTS

### Motor neuron- and interneuron-enriched slices show distinct patterns of gene expression

We exploited the dorsal to ventral anatomical differences in neuron classes within the spinal cord to create longitudinally sliced spinal cord samples relatively enriched for either motor neurons (MNslices; the ventral-most slice) or interneurons (INslices; the next more dorsal longitudinal slice) for our analyses. We chose this strategy for two reasons. First, while we are capable of performing single-cell qPCR analyses in spinal cord neurons, the technical limitations in working with single cells severely limited the number of gene products we could quantitatively screen from a single neuron, which would obviate our analysis of gene co-expression profiles. Second, by employing a pool of neurons from a slice in our analysis, we expect the heterogeneity to help “even out” the cell-to-cell variability in gene expression and allow for the major concerted and conserved changes in gene expression across cells to emerge as a signal above the “noise” of variability. Thus, with our slice samples we are likely detecting only the most profound and conserved changes in gene expression after SCI.

We compared RNA transcript numbers for 42 genes (see Appendix B) in longitudinal slices from the ventral surface of the lumbar spinal cord (L1-L5), where the Central Pattern Generator network driving hindlimb locomotion is predominantly located. Because motoneurons are located near the ventral surface of the cord, the first 300 μm slice in the grey matter is relatively enriched for motoneurons compared to the second, more dorsal slice, which is relatively more enriched for interneurons. Even though these slices enrich a given sample for a specific cell type, ultimately we are quantifying expression in very heterogeneous samples of spinal cord tissue. Therefore, it was possible that the mixture of multiple cell types in each sample, regardless of slice level, would simply occlude any differences in detectable gene expression between samples. This was not the case. In control animals, motor neuron-enriched slices (MNslices) showed very different expression patterns from interneuron-enriched slices (INslices). Figure 2A shows a cluster plot where each column represents a distinct gene, and relative expression is shown for ten MNslices and 9 INslices (rows) via a heat map based on the Z-score scaled within each gene. Hierarchical clustering cleanly subdivided the MNslices and INslices into two distinct nodes with 100% accuracy (Figure 2A). ANOVA revealed that 32 out of 42 genes were differentially expressed between MNslices and INslices (Figure 2A). The general trend was for greater expression of a given gene product in the INslices than the MNslices, with some exceptions: *GLRA1, CACNA1C, CHRNA2, SCN8A,* and *KCNA1* all showed higher expression in MNslices than INslices. These data demonstrate that differences in gene expression in specific neuron-enriched slices can be robustly detected even across very heterogeneous samples of spinal cord tissue.

**Figure 2.**
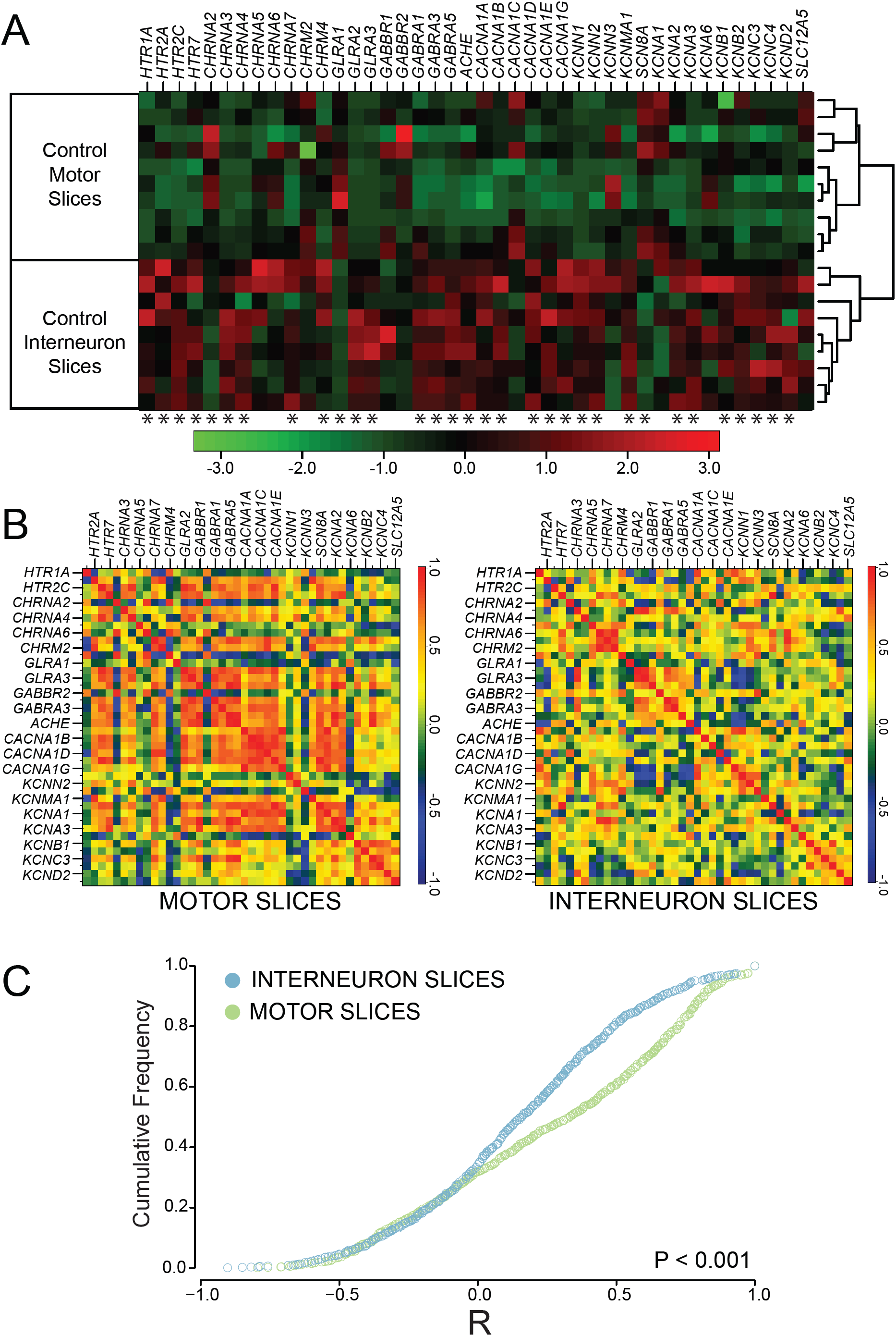
Channels and receptors are differentially expressed in interneuron- and motor neuron-enriched slices. **A)** Heat-map showing relative transcript levels of receptor and channel subtypes across different slice types of control (uninjured) animals. Data are expressed as a column Z-score where relative distance of a given expression value from the group mean is represented by color intensity. Each row is a different slice sample, which are clearly grouped into two distinct nodes by slice class as a result of post-hoc clustering. Genes were arbitrarily grouped based on functional subtypes of receptor and channel class. Asterisks under each column represent significantly different transcript levels between interneuron and motor neuron slices for a given gene (after Two-Way ANOVA, see Methods). **B)** Co-expression correlations among measured transcripts also differ between slice types. For each slice level from control animals, a correlogram was generated that displays the mean *R*-values for Pearson correlation tests as heat-mapped pixels for each pairwise comparison. Each *X*–*Y* coordinate represents one *R*-value for a given pairwise comparison. Along the diagonal is the autocorrelation for each gene, resulting in an *R*-value of 1.0 (red). Labels for every-other gene in the analysis are provided on the x- and y-axes for clarity, but both axes contain all genes used in the study in the same order. **C)** Cumulative distribution functions from the data shown in panel B reveals a significantly different distribution of R values, with higher co-expression in control MNslices (N = 10) than INslices (N = 9) between interneuron- and motor neuron enriched slices (D = 0.209, *P* < 0.001; Two-sample K-S test).

Absolute quantitation of mRNA abundance allows a level of analysis that goes beyond fold-change reporting of relative expression for a given gene. This analysis allows correlation-based approaches to provide a visualization of overall patterns of gene co-expression change across genes in different samples from a given experimental group (Garcia et al., 2014). In the case of these correlation analyses, the R-value for the correlation of expression of two genes across multiple sample measurements becomes the focal statistic parameter, as changes in the relationship between two genes’ expression levels can be determined both quantitatively (how “strong” is the correlation), as assessed by the magnitude of the R-value, and qualitatively (are two genes directly or inversely correlated) by the sign of the R-value. 42 genes of interest compared in a pairwise fashion resulted in 861 pairwise correlation coefficient results (R-values) for each experimental group. We describe the total population of R values for these 861 comparisons as a correlogram for each experimental group (Figure 2B), where R is heat mapped from red (R = +1.0) to blue (R = −1.0). Therefore, the strongest positive correlations for gene pair co-expression are seen as red squares, while those with weak or non-existent correlations are green, and those with a strong negative correlation are dark blue.

Using these analysis and visualization techniques, it is clear that the correlated patterns of gene expression differ between MNslices and INslices from control, non-injured animals. Cumulative distribution functions for R-values between INslices and MNslices (Figure 2C) reveal a significantly different distribution of correlations (two-sample K-S test, D = 0.209. *P* < 0.001). Under control conditions, INslices have a normal distribution of R-values (Figure 2C), representing a range of positive to negative correlations, with the median slightly above 0. In contrast, control MNslices have a more skewed R-value distribution (Figure 2C; median = 0.41) relative to the INslices (median = 0.16), with a fairly distinct set of positive correlations visible in this group (Figure 2B) Thus, expression of ion channel and receptor genes may be regulated in a more coordinated fashion in MNslices than in the INslices, or these analyses could reflect reduced neuronal heterogeneity in the MNslices.

### Spinal cord injury differentially affects gene expression in MNslices and INslices

To determine whether injury alters expression of channel and receptor genes differentially in MNslices and INslices, we performed a complete spinal cord transection at spinal level T8-9. After 30-60 days, the lumbar spinal cord was dissected, sliced in 300 μm longitudinal sections, and prepared for analysis as above (Figure 1). We quantified 42 different transcripts of voltage-gated ion channels and for acetylcholine, GABA, glycine, and serotonin receptor subtypes. Combined channel and receptor expression patterns analyzed via 2-way hierarchical cluster analysis based on Pearson distance (Figure 3) divided the population of 37 different slice samples into distinct groups that correspond with almost 100% fidelity to control MNslices, SCI MNslices, and INslices (combined). The first clustering distinction is made with extremely high fidelity between INslices and MNslices, demonstrating that these two anatomical slice level distinctions have robust and distinct patterns of gene expression, as was seen in Figure 2. The node containing the MNslices subdivides cleanly into two groups, corresponding to SCI and control samples (Figure 3), showing that the pattern of gene expression changes markedly after SCI (see below). The node containing the INslices subdivides into distinct control and SCI groups (Figure 3), as well, although with somewhat less homogeneity than the motor neuron slices. The control INslices cluster as one distinct group, but the SCI INslices are contained within two distinct nodes in the analysis. Nevertheless, there is a striking amount of fidelity of clustering with the experimental levels of injury and slice level: only one slice did not cluster with the appropriate experimental group: a Control INslice (ConInt7) clustered with the Control MNslices (Figure 3, see asterisk). These data demonstrate clear distinctions in expression profiles associated with spinal cord injury in these groups.

**Figure 3.**
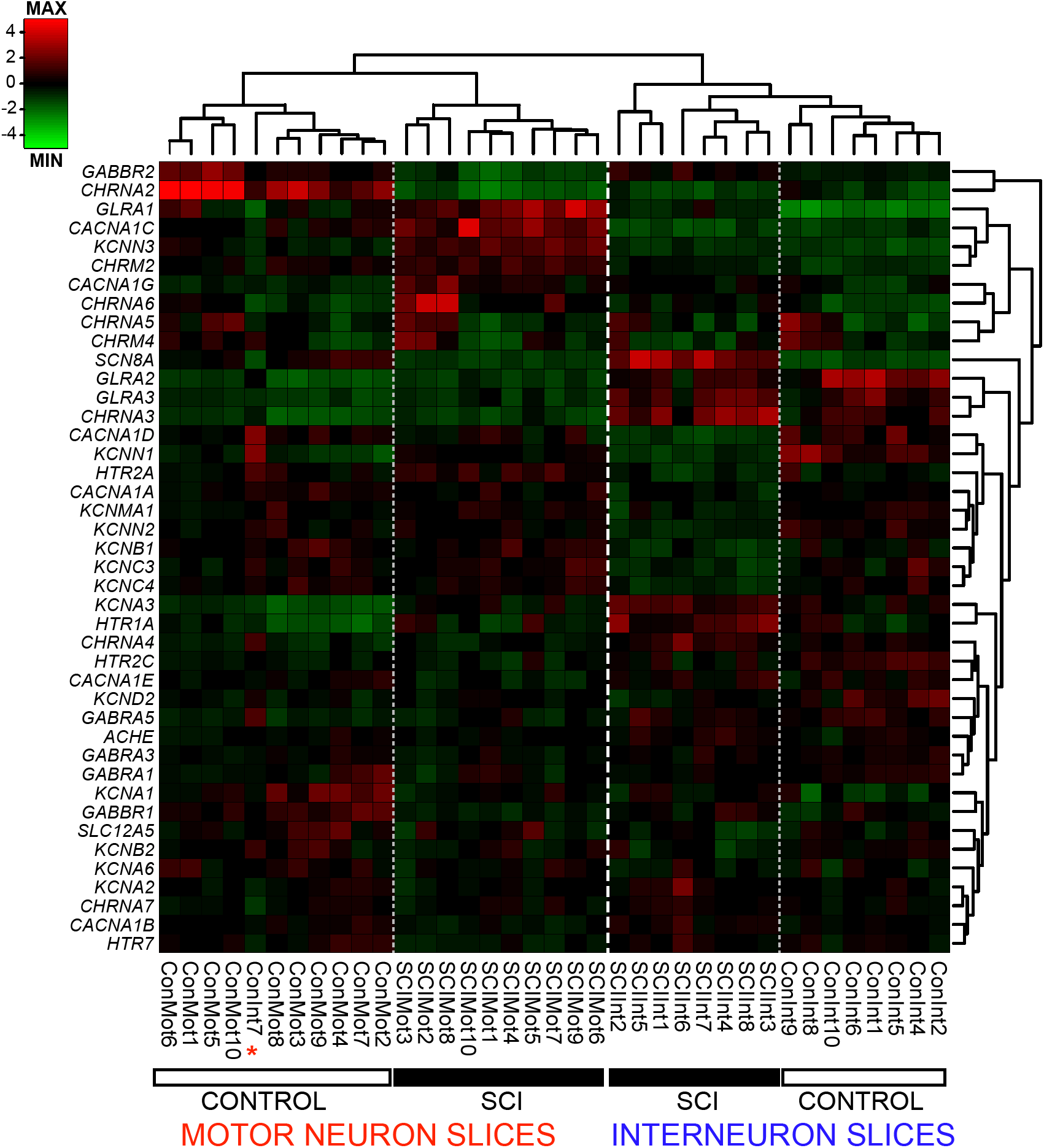
Ion channel and receptor transcript profiles correspond to slice level and injury state in lumbar spinal cord. Dendrograms and heat map of individual slice samples (columns) from control and SCI motor- and interneuron-containing slices for channel and receptor transcript levels (rows). Hierarchical clustering of sorted slice expression profiles distinctly associated samples from each of the four groups, with one exception (red asterisk). Con = control, SCI = injured, Mot = motor neuron-enriched slices, Int = interneuron-enriched slices. Relative expression is indicated by Z-score based on the average transcript levels for each gene (rows), and not each sample (columns).

### Ion channel and receptor genes show widespread changes in expression following SCI in both MNslices and INslices

We specifically focused on subsets of ion channel and receptor genes for this study to start to understand the underlying mechanisms for changes in excitability and transmitter sensitivity that have been reported in spinal neurons following SCI (e.g. Li et al. 2004; Murray et al. 2011; Husch et al. 2012). There was widespread change in ion channel expression levels in both MNslices and INslices following SCI. Of 20 ion channel genes studied, only 4 (*CACNA1, CACNA1E, KCNA6,* and *KCNB2)* showed no changes in either MNslices or INslices (Figure 4). There were also no differences in the expression of the chloride transporter *SLC12A5* in our slices after injury. Eight of 16 channel genes changed in both MNslices and INslices; of the remaining 8 genes, 4 changed only in INslices and 4 changed only in MNslices (Figure 4). In MNslices, 11 out of 12 genes that changed *increased* significantly in expression levels after SCI while only 1 (*SCN8A*) decreased in expression. Conversely, in INslices 7 genes decreased in their expression levels while 5 genes were upregulated. In most instances (7 out of 8 times) when changes were found in both MNslices and INslices for a particular channel gene, they changed in opposite directions: most commonly a given channel was downregulated in INslices but upregulated in MNslices (Figure 4). There were no changes detected in expression of the *SLC12A5* chloride transporter.

**Figure 4.**
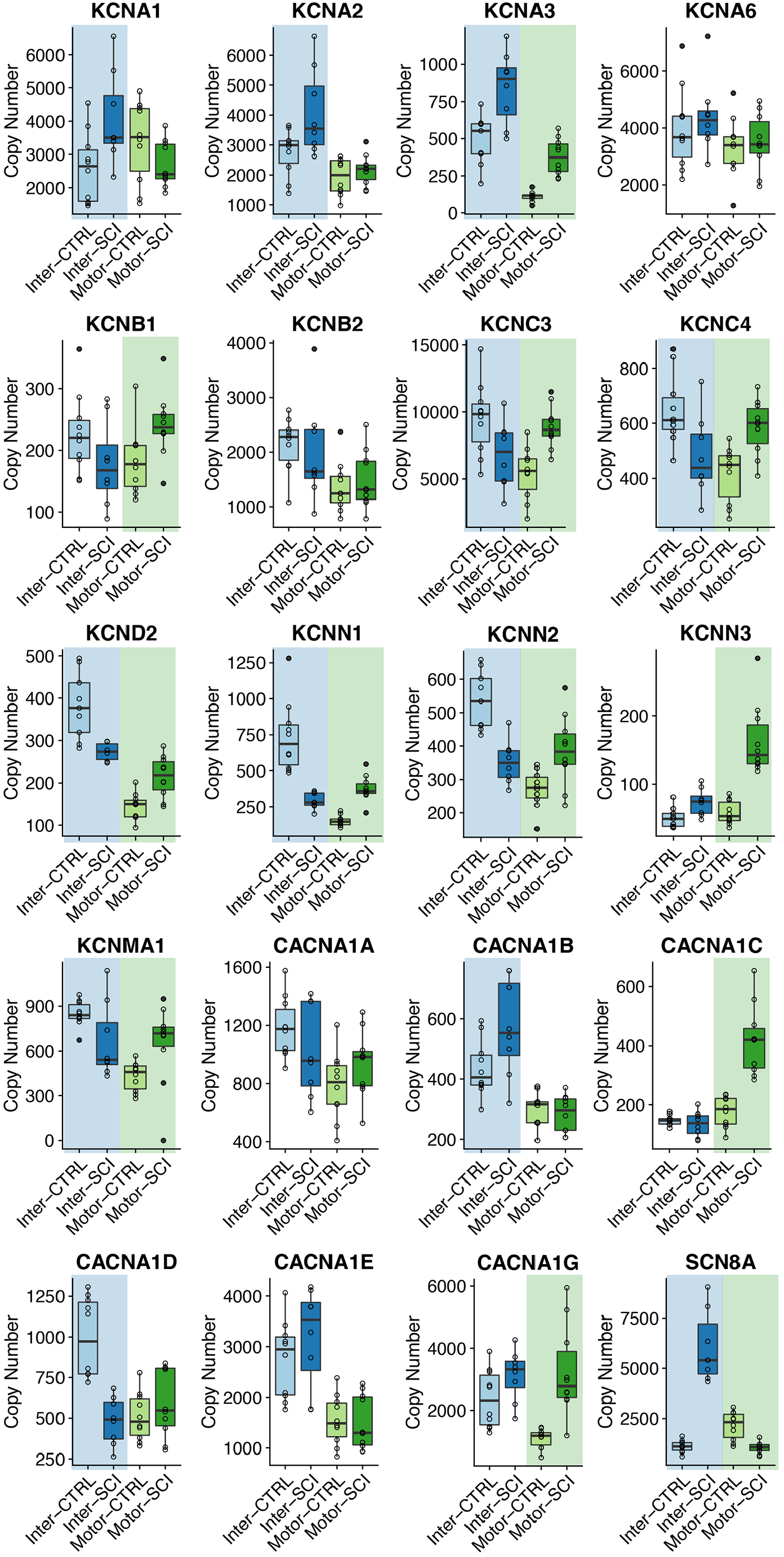
Boxplots for mRNA copy numbers for each ion channel gene of interest across all four experimental groups. INSlices are represented in shades of blue, while MNSlices are represented in shades of green. For a given boxplot, the median is denoted by a horizontal line, and the box extends to the 25^th^ and 75^th^ percentiles. Individual observations are presented as open circles, and whiskers extend to the most extreme values that are within the interquartile range. Outliers are defined as points outside 1.5 times the interquartile range above the upper quartile and below the lower quartile, and designated by filled circles. Significant differences (p < 0.05; post-hoc *t*-test following Two-Way ANOVA) between control and injured samples for a given slice level are denoted with a shaded background of the appropriate color (blue or green).

Changes in receptor expression were less prevalent than for ion channels (Figure 5). While 16/20 channel genes showed a change of expression in either the INslice or MNslices after injury, only 11/21 receptor genes showed a significant change. Of those 11 genes, 8 receptor types changed in INslices and 7 changed in MNslices. All cases of significant changes in receptor expression, whether in MNslices or INslices, were *increases* in expression in SCI individuals with two exceptions: *GABBR2* was downregulated in MNslices in SCI animals, while CHRNA2 was almost undetectable in MNslices after SCI. Four of the 11 receptor genes that changed expression levels did so in both slice types, while 7 genes changed in only one slice type (3 MNslices and 4 in INslices [Figure 5]). Acetylcholinesterase expression (*ACHE*) was significantly upregulated in SCI INslices, but no differences were detected in MNslices. These patterns of expression demonstrate not only that channel and receptor gene expression are influenced widely after SCI, but that there is not a simple unidirectional change in expression. Rather, patterns of expression changes are distinct both in increasing or decreasing across gene types, as well as changing in distinct patterns between slice levels.

**Figure 5.**
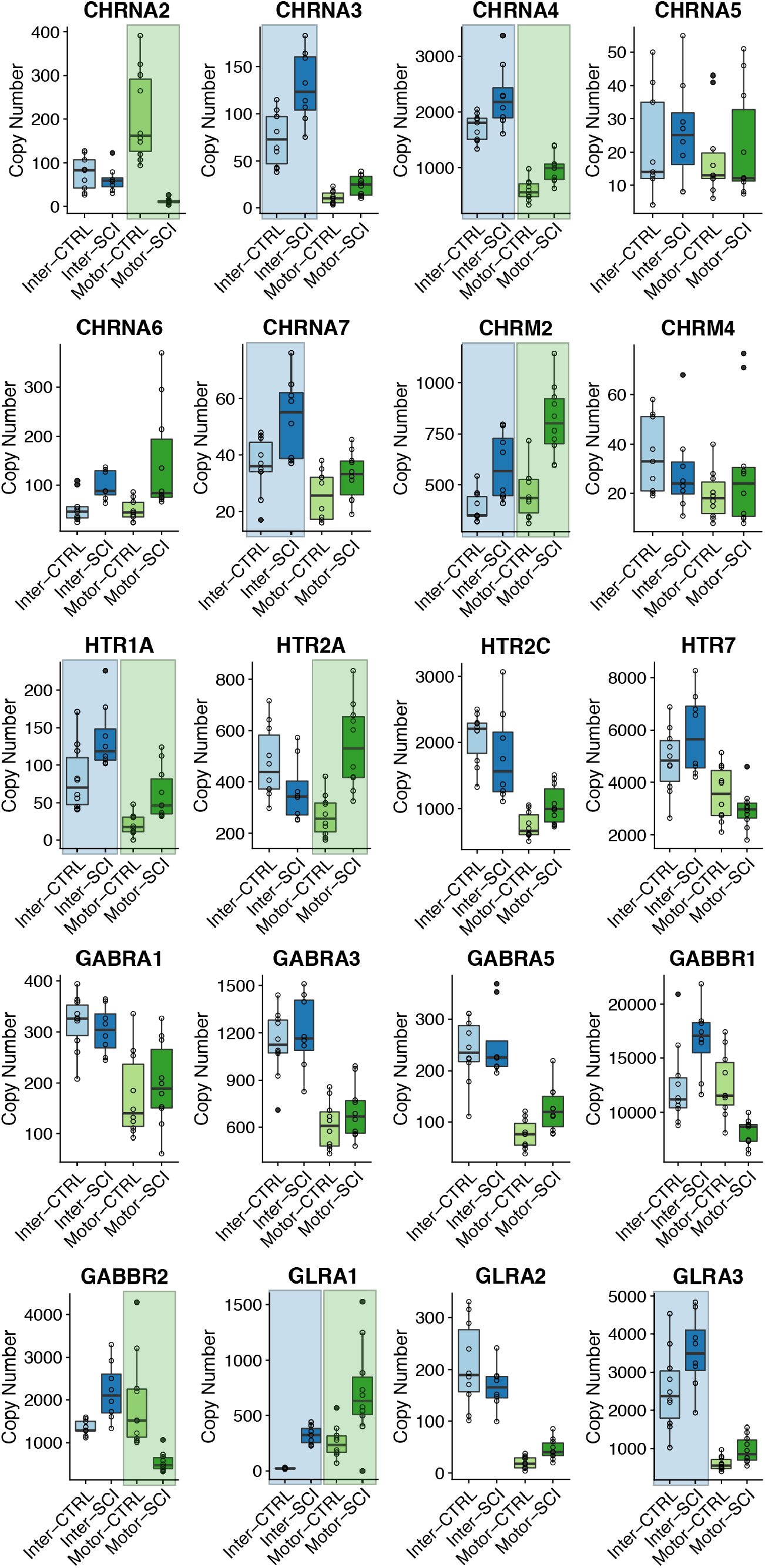
Boxplots for mRNA copy numbers for each receptor gene of interest across all four experimental groups (motor and interneuron slices, control and injured). Formatting of groups, boxplots, and statistics as in Figure 4.

### Changes in ion channels and receptors at the MNslice level are consistent with changes seen at the level of individual motor neurons

Although the longitudinal slices will enrich each sample for a particular spinal cord neuron subtype, the slice samples still represent thousands of cells in a very heterogeneous mix. Unfortunately, our quantitative measurements of gene expression were not sufficiently sensitive to quantitate multiple gene transcripts from singly collected motor neuron samples, and so we were limited in the number of transcripts that we could compare. To determine whether the enriched slices were representative of individual cellular expression changes, we compared the MNslices expression profiles to isolated motor neuron samples for 4 target genes: *CACNA1A, CACNA1D, HTR2C,* and *SCN8A*. For all four genes, the same expression pattern was seen at the single cell and slice levels: *SCNA8A* was lower in motor neurons from SCI animals, *HTR2C* expression was higher in motor neurons from SCI animals, and *CACNA1A* and *CACNA1D* levels were not different between control and SCI motor neurons (Figure 6). Our previous analysis of increases in HTR2c expression after SCI did not quite reach statistical significance for MNslices (P = 0.092; Figure 5), due to the extensive multiple comparison adjustments that were necessary for pairwise *t-*tests. However, in our simpler comparison in Figure 5, *HTr2C* mRNA levels were found to be significantly higher in both MNslices and motor neuron samples (Figure 6). While we could not validate gene expression for every gene in both general cell/slice types, these data provide evidence that expression changes at the longitudinal slice level are reflective of changes occurring in individual neurons of a given type.

**Figure 6.**
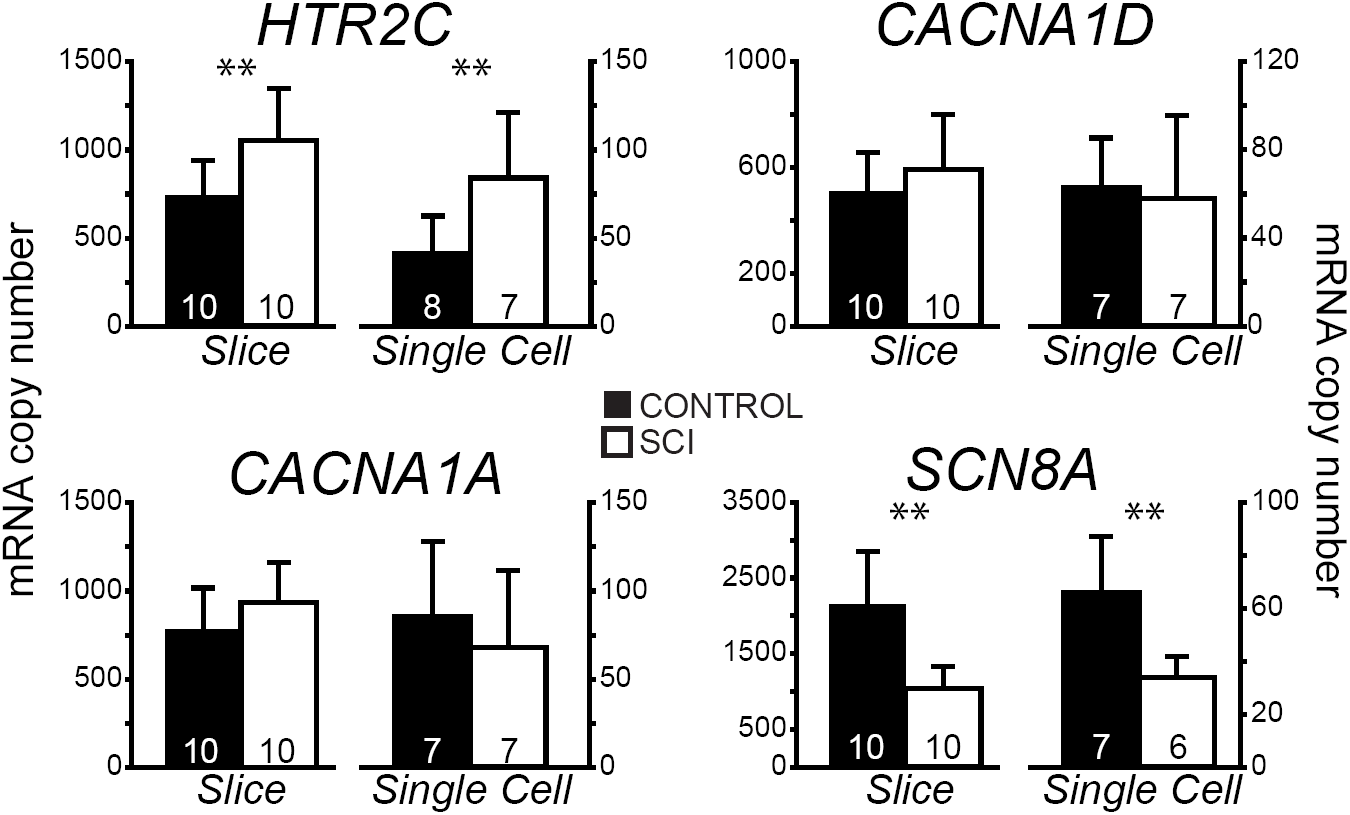
Single motor neuron gene expression analyses recapitulate transcript levels in whole slices. Mean ± SD copy numbers of four genes of interest across experimental groups (control and injured) in both slices and single motor neurons. Significant differences (p < 0.05; *t*-test) between control and injured samples are shown with **. Note, while our post-hoc analyses following two-way ANOVA did not have the statistical power to detect significant differences in HTR2C in motor neuron slices (see Figure 5), a direct comparison with *t*-test reveals this difference in both slices and single motor neurons. Sample sizes indicated in each bar.

### Patterns of correlated gene expression change are different in MNslices and INslices after SCI

We generated correlograms (as in Figure 2B) from both control and SCI animals for MNslices (Figure 7A1, 7B1) and INslices (Figure 8A1, 8B1) to look for changes in the pattern of co-expression of all gene pairs. Beside each correlogram is a histogram showing the frequency distribution of R-values for the population of correlation coefficients in that experimental group (Figure 7A2 and 7B2, Figure 8A2 and 8B2). We also generated gene co-expression network plots for all of the groups analyzed (Figure 7A3 and 7B3; Figure 8A3 and 8B3), where each correlated pair of transcripts (R > 0.7) is connected with a line to visualize the density of correlated mRNAs for a given experimental group. Finally, the change in correlation profiles as a result of injury is shown for MNslices (Fig. 9A1) and INslices (Fig.9B1) as the difference in R-values (R_SCI_ - R_CONTROL_), and plotted as ΔR in equivalent fashion as the raw R-values. In these ΔR plots, injury-induced increase in correlation between expression of 2 genes is shown in yellow and red, no change is shown in green, and injury-induced decreases in co-expression are shown in blue. These changes can also be visualized both by plotting the frequency distribution of these ΔR values (Figures 9A2 and 9B2) as well as the cumulative distribution of R-values compared between Control and SCI slices (Figures 9A3 and 9B3).

**Figure 7.**
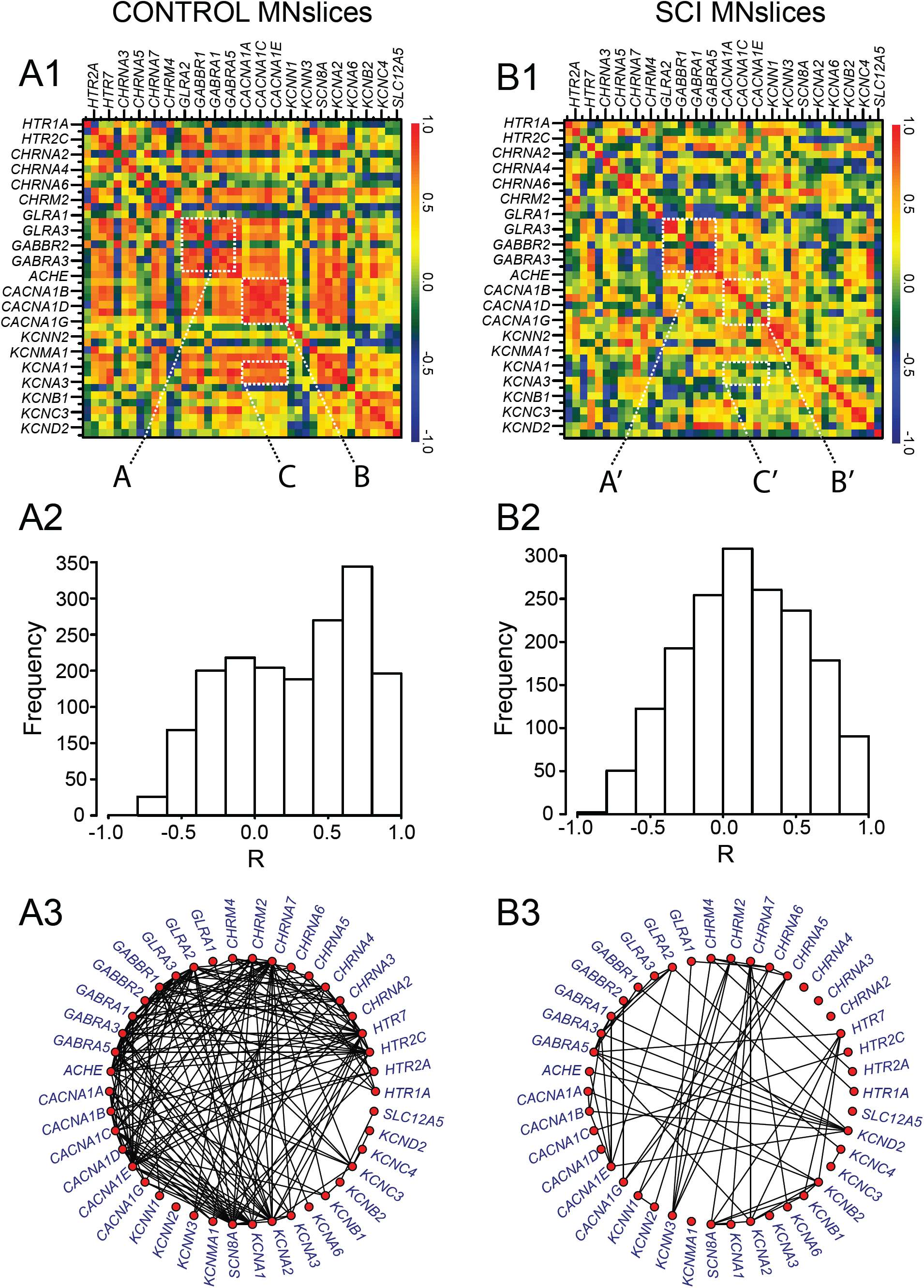
Changes in correlated gene expression as a result of SCI in MNslices. **A1 and B1)** Correlograms were generated as described in Figure 2. The control correlogram is re-plotted from Figure 2 for comparison with the SCI correlogram below. Dashed boxes are labeled (e.g. A, A’) for discussion of Results in the main text. The order of the genes along the x- and y-axes are the same, and are represented in the same order as in Appendix B for comparison. **A2 and B2)** The histogram for the distribution of R-values in a given experimental group is plotted beside each correlogram. **A3 and B3)** Gene co-expression network plots were generated to visualize the density of correlated transcript pairs before and after SCI. Each red node represents a given transcript, and if two nodes are connected with a line, their mRNA levels were correlated with an R-value > 0.7.

**Figure 8.**
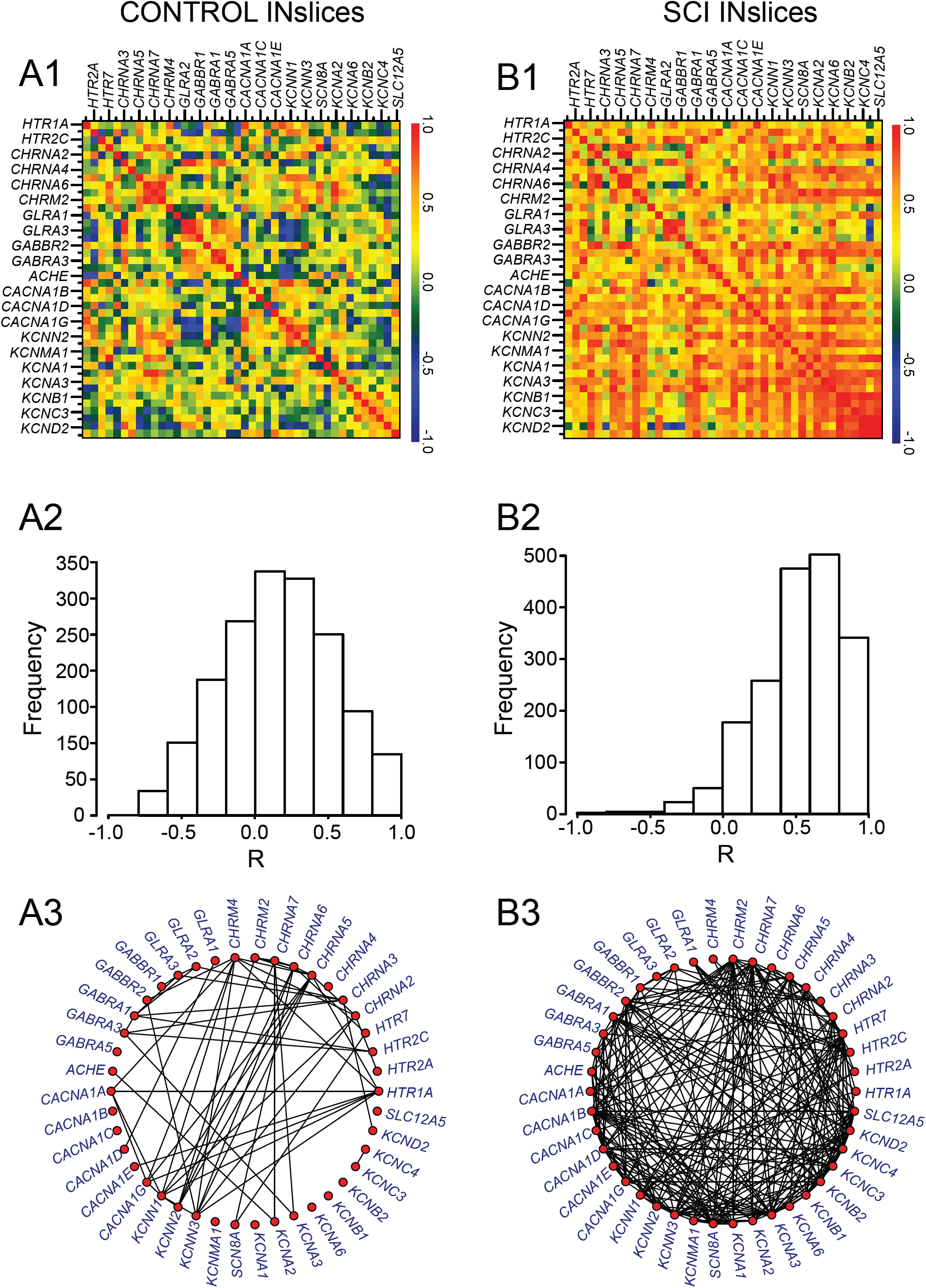
Changes in correlated gene expression as a result of SCI in INslices. All notation as in Figure 7. Note the substantial number of connected nodes in panels **8A3 vs. 8B3** that reflect an increase in positively correlated gene co-expression in SCI as a reflection of the histograms in panels **A2 and B2.**

**Figure 9.**
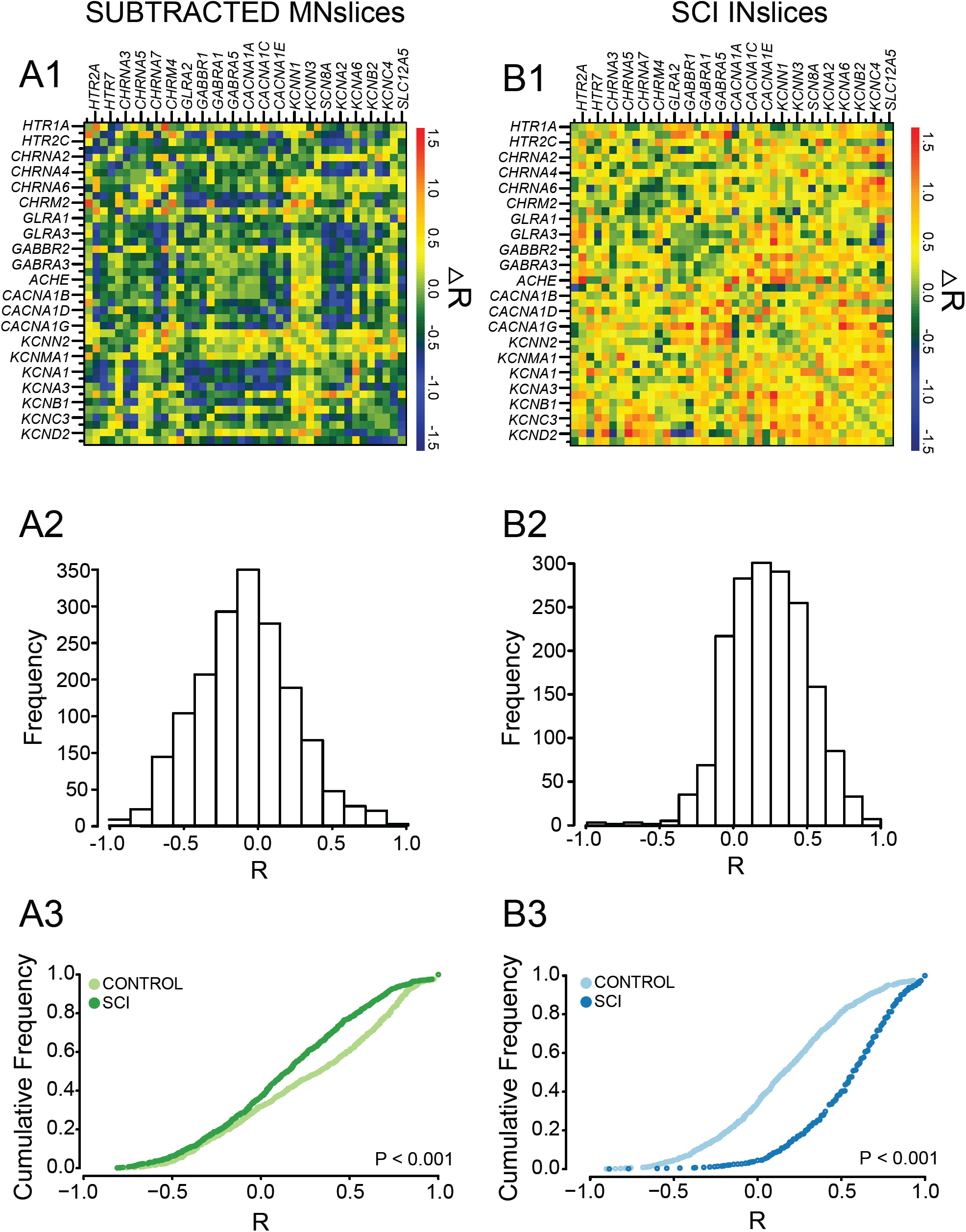
Correlation analyses reveal that interneuron-enriched slices and motor neuron-enriched slices respond differently to spinal cord injury. **A1 and B1)** Correlograms were generated (as described in Figure 2) for subtracted R-values (ΔR = R_SCI_ – R_CTRL_) for each transcript pair to detect changes in relationships among transcripts as a result of injury. **A2 and B2)** The histogram for the distribution of ΔR in a given experimental group is plotted below each correlogram of ΔR-values. **A3 and B3)** Cumulative distribution functions from each slice type reveals a significantly different distribution of R values (*P* < 0.001; Two-sample K-S test) between control and SCI animals. Motor neuron-enriched slices tend towards a loss of positive correlations, as seen by the leftward shift of the distribution for values greater than 0 from control to SCI. Conversely, interneuron-enriched slices show a dramatic rightward shift overall, as a manifestation of the large number of relationships that become positively correlated following SCI as shown in the panels above.

Using these combined analyses and visualization techniques, two major conclusions can be drawn. First, it is clear that the correlated patterns of gene expression change after SCI in both MNslices and INslices (Figures 7 and 8). Second, the changes in correlations are very different for MNslices and INslices. In the SCI animals, MNslices show a distinct reduction in overall correlation of expression: the R-value distribution changes from being skewed toward positive values in controls to a more normal distribution with a median closer to 0 after SCI. This could be due to changes that either increase or decrease the level of any given transcript – ultimately altering the relationships among transcript levels. This is borne out in the ΔR distribution for MNslices, where there is a shift towards a negative ΔR (Figure 9A2) with a median value of ΔR = −0.127 that reflects the overall tendency towards loss of correlation after SCI. Furthermore, the gene co-expression network plots clearly reveal a substantial decrease in the density of connections between transcripts in the SCI MNsclices relative to the Control MNslices (Figure 7A3 vs. 7B3). Some genes completely lose their co-expression relationships with the other measured genes, such as the cholinergic receptor genes CHRNA2-4, while the large majority of genes showed a marked reduction in overall connections. Only a few, including several voltage-dependent potassium channels in the KCN family, show net increases in numbers of connections. This overall net decrease in co-expression is demonstrated by a significant decrease in median R-value in MNslices following SCI (Figure 9A3). We see exactly the opposite trend in INslices (Figure 8). INslices change from a normal distribution of R-values in control animals to a highly skewed distribution of much more positive R-values in SCI (Figure 8A2 and 8B2), indicating a large-scale *increase* or *gain* of positive correlations in gene expression after SCI; again, this could arise either through increased or decreased mRNA levels for any given transcript. This is further demonstrated by a positive ΔR in INslices after SCI with a median value of ΔR = 0.367 (Figure 9B2), a significant increase in median R-value distribution in INslices after SCI (Figure 9B3), and a substantial increase in density of connections in the gene co-expression network plot (Figure 8A3 vs. 8B3). After SCI, every gene had at least one significant co-expression with another gene, and virtually all the genes increased their number of connections.

It is also possible using this visualization procedure to identify distinct correlation relationships among sets of genes within a slice level that have different fates after SCI. First, in MNslices there is a clear positive or negative correlation of expression among glycine and GABA receptor subtypes (Figure 7A1, *box A*), and these correlations largely persist in SCI animals (Figure 7B1, *box A’*). In a contrasting example, there are positive correlation relationships that can be seen among voltage-gated calcium channel genes (*CACNA1A-1G*; Figure 7A1, *box B*) and between calcium channel and voltage-gated shaker-related K+ channels of the *KCNA* family (Figure 7A1, *box C*) in MNslices of control animals. These correlations are lost or substantially weakened in SCI animals (Figure 7B1, *boxes B’* and *C’*), resulting in ΔR-values in the subtracted correlogram that color code blue as lost correlations (Figure 9A1). These results demonstrate that different subsets of genes undergo differential regulation after SCI in MNslices. In stark contrast, in INslices almost all relationships among genes shift towards a positive correlation (Figures 8 and 9). This shows a striking uniformity of shift in expression patterns of channels and receptors in INslices of SCI animals, and underscores how different responses can be between distinct slice levels even in the same population of individuals.

## DISCUSSION

Our results demonstrate that SCI does not result in uniform changes in gene expression within the lumbar spinal cord: different longitudinal slices associated with relative enrichment for motor neuron and interneuron pools experience substantially different changes in expression in response to injury. In comparing MNslices and INslices, we can clearly see that different classes of neurons within the spinal cord have distinct expression profiles, both in control and injured conditions. This is demonstrated first in our clustering analyses: samples cluster first and foremost by slice type (MNslice vs. INslice). Then within a slice type, the samples cluster convincingly into control and SCI states. The difference between slice types is perhaps best illustrated by the fact that for a given gene that was differentially expressed in control and SCI animals, in 7 of 8 cases the changes in abundance were in *opposite* directions in MNslices and INslices. Thus, while it is perhaps not surprising that different regions of the cord react differently to SCI, this clear evidence underscores the importance of recognizing there is a not a uniform change in the cord after an injury.

Second, we see widespread changes in relative abundance of a given transcript following SCI. It is clear that SCI causes widespread changes in gene expression in the spinal cord across numerous gene families: this has been demonstrated through the use of qPCR (Esmaeili and Zaker, 2011; Di Narzo et al., 2015), microarrays (Carmel et al., 2001; Ryge et al., 2008, 2010; Wienecke et al., 2010; Liu et al., 2014) and RNAseq (Chen et al., 2013; Lee-Liu et al., 2014). Our results add to this body of work, and can help provide insight into potential mechanisms underlying changes in physiology following SCI. One of the most common physiological impacts of SCI is on motor neurons below the site of injury which show hyperexcitability, ultimately influencing spasticity in affected limbs (Bennett et al., 2001; Raineteau and Schwab 2001; Adams and Hicks 2005). This spasticity is associated with physiological changes that are mirrored in our expression results. For example, spasticity in sacral motor neurons in chronically spinalized rats is associated with changes in both sodium- and calcium-mediated persistent inward currents (Li et al., 2004). This is in part due to upregulation of Ca_v_1.2 alpha-subunits, while Ca_v_1.3 remained unchanged (Anelli et al., 2007). This is a direct parallel with our results: *CACNA1C* (Ca_v_1.2) was significantly higher in MNslices of chronic SCI, while *CACNA1D* (Ca_v_1.3) levels were not different. Modeling work also suggests a potential interaction between these persistent currents and calcium-activated potassium (KCa) currents in hyperexcitability after SCI (Venugopal et al., 2012), and we saw significant changes in 3 SKKCa (*KCNN1,2,3)* and a BKKCa (*KCNMA1*) subunits in chronic injured MNslices. Muscle spasms after chronic SCI have also been associated with enhanced serotonin receptor activity, including 5-HT2B and 5-HT2C receptors in motor neurons (Murray et al., 2011). These are also consistent with our data, as we see increases in *HTR2A* and *HTR2C* serotonin receptor mRNAs in MNslices. Finally, one of the most striking expression changes we detected was the almost complete loss of *CHRNA2* expression, known to encode the nicotinic receptor alpha-2 subunit, in MNslices. *CHRNA2* is selectively expressed in Renshaw cells of the ventral spinal cord (Perry et al., 2015), where they receive cholinergic input from alpha motor neurons as part of the recurrent inhibitory feedback circuit of motor neuron excitability (Moore et al., 2015). Loss of this connection between alpha motor neurons and Renshaw cells, which ultimately feed back to inhibit motor neurons, could be a novel mechanism involved in spasticity following SCI as well as other motor neuron diseases such as ALS (Wootz et al., 2013).

Third, our data show that changes in gene expression after SCI do not simply alter the abundance of a given gene product, but also substantially change co-expression relationships *across* genes resulting in changes in correlated levels of transcripts. Ion channels and receptors work in concert to adjust the excitability and firing pattern of a given neuron. It is therefore imperative to expand gene expression profiling to consider the relationships across genes which participate in co-expression networks (Schulz et al., 2007; Hawrylycz et al., 2011; Menashe et al., 2013; Garcia et al., 2014; Grange et al., 2014). For example, GABA and glycine mediate the two most important inhibitory synaptic systems in the spinal cord (Schneider and Fyffe, 1992; Kiehn, 2006), and previous work has identified co-localization of their receptors (Todd and Sullivan, 1990; Todd et al., 1996). Our data identify a positive correlation among GABA and glycine receptor subunit mRNA levels, particularly in MNslices, supporting the identified action of GABA and glycine as co-transmitters in premotor interneurons (Jonas, 1998; O’Brien and Berger, 1999), and this co-expression is resistant to SCI in MNslices. These results suggest that while disruption in GABA/glycine signaling in the dorsal spinal cord may be an important underlying cause of hyperexcitability underlying neuropathic pain in spinal cord injury (Gwak and Hulsebosch, 2011), there may not be a major effect of SCI on expression of genes associated with inhibitory synaptic transmission in the ventral cord.

Another example of changes in expression as a result of SCI is seen in channel relationships in MNslices among voltage-gated calcium channels (*CACNA1A-G*) and potassium channels of the shaker-related family (*KCNA1-3*). In control animals, there is positively correlated expression among many of these calcium and potassium channels. Following SCI there is a widespread disruption of this correlated expression, even though mean RNA levels change significantly in only 3 of these 9 genes in MNslices. This could be due to either an increase or decrease in expression of a given transcript, or both an increase and decrease in transcripts involved in a correlation. One cellular mechanism to regulate bursting activity in motor neurons is the relative ratio of voltage-gated calcium and A-type potassium conductances (Ball et al., 2010; Franklin et al., 2010; Hudson and Prinz, 2010). By altering this ratio, motor neuron intrinsic activity could be substantially altered, which is consistent with reported changes in Ca^2+^ and K^+^ currents in SCI neurons (Nashmi et al., 2000; Nashmi, 2001; Li et al., 2004; Anelli et al., 2007; Venugopal et al., 2012). The molecular data presented in our results, while inferential, could therefore shed light on potential mechanisms underlying excitability dysfunction in motor neurons after SCI.

Fourth, our results demonstrate that changes in co-expression patterns after SCI vary profoundly in different regions of the spinal cord. MNslices tend to experience a net loss of gene correlation among potentially interacting channel and receptor proteins, while INslices show a massive increase in the number of correlated genes across many receptor and channel types. Activity changes as a result of loss of descending projections have been shown to cause decorrelation in channel expression in invertebrate motor neurons (Temporal et al., 2012, 2014), especially between voltage-gated Ca^2+^ and K^+^ channels (Temporal et al., 2014), and our data are consistent with this effect in vertebrate MNslices. Perhaps the most striking change in our results, however, is the widespread increase of channel and receptor co-expression after SCI in INslices. We can only speculate about the implications or mechanisms underlying these changes, as nothing of this nature has previously been reported. However, these results are reminiscent of differences in channel and receptor co-expression in mouse spinal cord across postnatal development (Garcia et al., 2014). Young pups show widespread correlated patterns of channel and receptor expression in their cords that over the course of postnatal development get pared down to specific subsets of correlated genes (Garcia et al., 2014). Therefore, it is possible – although pure speculation – that loss of descending tracts to the lumbar spinal cord as a result of injury could induce a developmental “reversion” of sorts in interneuron populations. Regardless of whether this hypothesis is on the right track, the primary implication of our data is that neurons in distinct cell regions of the lumbar cord respond substantially differently with respect to expression changes after SCI.

These widespread changes in gene expression not only have implications for physiological impacts of SCI, but also demonstrate that SCI has fundamental impacts on overall transcriptional regulation processes. The most obvious potential mechanism for changes in co-expression of multiple genes is alterations in transcription factors associated with regulating production of the mRNAs we measured. SCI is known to affect many transcripts (Ryge et al., 2010; Wienecke et al., 2010), and identifiable gene clusters emerge with distinct temporal dynamics after SCI; each of these clusters can be attributed to differentially expressed transcription factor subtypes (Ryge et al., 2010; Zhang and Wang, 2016). These kinds of gene clusters and their associated regulatory factors play a major role in the establishment of gene co-expression networks in developing spinal cord (Jessell, 2000; Lee and Pfaff, 2001), and are likely targets for genetic impacts of SCI. However, other mechanisms for changes in steady-state mRNA levels have also been associated with SCI, such as alterations in microRNA expression (Yunta et al., 2012; Nieto-Diaz et al., 2014). It is likely that multiple transcriptional and post-transcriptional mechanisms will be evoked as a result of such widespread and traumatic changes as caused by SCI.

Taken together, our results shed greater light on the complex relationships in gene expression in the spinal cord, and the differential effects of injury on these relationships both across neuron classes and gene families. However, there are serious limitations to the analysis at hand. First, mRNA does not represent mature and functional protein, and so at best the link between our data and physiology is inferential. Second, while our technique of longitudinal slices uses the anatomical features of the spinal cord to our advantage, we are still pooling heterogeneous groups of neurons and glia together for these analyses. This is especially true in the so-called “interneuron slices” where we have combined multiple spinal laminae that serve diverse systems as their final targets. Therefore, while we have likely detected concerted changes in gene expression above this “noise,” it is also possible that we have failed to detect meaningful physiological changes as a result. Careful follow-up studies are required to provide any functional insight into the consequences of these changes in gene expression. In particular, future work will need to control for more cell-type specificity, as well as distinctions between flexor and extensor CPGs, to provide more meaningful mechanistic insight. However, we are optimistic that these data can help to better target potential physiological mechanisms for more specific, cellular level experiments that can enhance the resolution of our findings as well as provide direct mechanistic insight. With target channel and receptor targets identified *a priori*, we are hopeful more efficient experimental insights can be gained.

## ACKNOWLEDGEMENTS

This work was funded by a grant from the Missouri Spinal Cord Injuries Research Program (DJS), Craig H. Neilsen Foundation (DJS), and NIH grant RO1-NS17323 (RHW). We thank Benjamin McGlaughon for technical assistance in the experiments and the behavioral analysis of functional recovery after spinal cord injury.

